# Repurposing non-cancer Drugs in Oncology – How many drugs are out there?

**DOI:** 10.1101/197434

**Authors:** Pan Pantziarka, Vidula Sukhatme, Lydie Meheus, Vikas Sukhatme, Gauthier Bouche

## Abstract

**Background:** Drug repurposing can speed up access to new therapeutic options for cancer patients. With more than 2000 drugs approved worldwide and 6 relevant targets per drug on average, the potential is quantitatively important. In this paper, we have attempted to quantify the number of non-cancer drugs supported by either preclinical or clinical cancer data.

**Methods:** A PubMed search was performed to identify non-cancer drugs which could be repurposed in one or more cancer types. Drugs needed at least one peer-reviewed article showing an anticancer effect in vitro, in vivo or in humans.

**Results:** A total of 235 eligible non-cancer drugs were identified (Table 1). Main charateirstics of the drugs are summarized in Table 2. 67 (29%) are on the WHO list of essential medicines and 176 (75%) are off-patent. 133 (57%) had human data in cancer patient(s). Four were listed in clinical guidelines, namely thalidomide, all-trans retinoic acid, zoledronic acid and non-steroidal anti-inflammatory drugs (NSAID). Several drugs have shown a survival benefit in randomized trials such as cimetidine (colorectal cancer), progesterone (breast cancer) or itraconazole (lung cancer). Several other drugs induced responses in rare tumours, like clarithromycin, timolol or propranolol.

**Conclusion:** We have found that the number of off-patent repurposing opportunities is large and increasing. Joint non-commercial clinical development (academics, governments, charities) may bring new therapeutic options to patients at low cost, especially in indications for which the industry has no incentive to invest in.

## Introduction

Drug repurposing can speed up access to new therapeutic options for cancer patients. Whereas it is not unusual to attempt to find new cancer uses for existing anticancer drugs, less attention and efforts are made to find anticancer uses of non-cancer drugs ^1^. However, many non-cancer drugs could be potentially repurposed against cancer though to our knowledge no figures have been put forward so far ^2^. One advantage of non-cancer drugs is that they represent a way to adapt to new knowledge about cancer. For instance, Tadalafil (PDE-5 inhibitor, erectile dysfunction), inhibits myeloid-derived suppressor cells (MDSC) in cancer patients ^3^. Or propranolol (beta-blocker, hypertension) reduces proliferation and migration of angiosarcoma models ^4,5^, by blocking beta-adrenergic receptors expressed by angiosarcoma cells ^6^.

With more than 2000 drugs approved worldwide and 6 relevant targets per drug on average ^7^, the potential is quantitatively important. What’s more, with regular new drug approvals, the toolbox that repurposed drugs represent is growing every year. Even antibodies are now being repurposed (e.g. rituximab in pemphigus ^8^), sometimes based on the discovery of off-target effects ^9,10^.

When a drug loses its patent protection, the incentives for the market authorization holder are also lost. Sometimes, other private entities attempt to protect the new therapeutic use by various means and undertake a commercial development^11^. However, this strategy remains financially risky, which makes it less attractive to investors and venture capitalists compared to other biotech investments. We have called these drugs “financial orphan drugs” ^12^.

In this paper, we have attempted to quantify the number of non-cancer drugs supported by either preclinical or clinical cancer data.

## Methods

A PubMed search was performed to identify non-cancer drugs which could be repurposed in one or more cancer types. Drugs needed at least one peer-reviewed article showing an anticancer effect in vitro, in vivo or in humans.

## Results

A total of 235 eligible non-cancer drugs were identified (Table 1). Main characteristics of the drugs are summarized in Table 2. 67 (29%) are on the WHO list of essential medicines and 176 (75%) are off-patent. 133 (57%) had human data in cancer patient(s). Four were listed in clinical guidelines, namely thalidomide, *all-trans* retinoic acid, zoledronic acid and non-steroidal anti-inflammatory drugs (NSAID). In the first 3 cases, pharmaceutical companies took the lead and re-branded or re-formulated the drugs. This was not the case for NSAIDs, listed in desmoid tumours guidelines and used off-label.

**Table 1:** List of non-cancer drugs with at least one peer-reviewed paper supporting its use against cancer.

| Drug | Pharmacological class | Main Indications |
| --- | --- | --- |
| Acetazolamide | Carbonic anhydrase inhibitors - ATC Code: S01EC01 | Glaucoma, diuretic, epilepsy |
| Acetaminophen (paracetamol) | NA | Analgesic |
| Agomelatine | Melatonergic antidepressant | Insomnia |
| Albendazole | NA | Anthelmintic |
| Aliskiren | Direct renin inhibitor | Essential hypertension |
| Allopurinol | Xanthine oxidase inhibitors | Gout |
| All-trans retinoic acid (tretinoin) | NA | Acne, APL |
| Alpha-Lipoic Acid | Thioctic acid - A16AX01 | Diabetic neuropathy (Germany) |
| Amantadine | Anticholinergic | Parkinson, Influenza A |
| Amiloride | Potassium-sparing diuretic | In congestive heart failure or hypertension treated with thiazides, to conserve potassium |
| Amiodarone | Antiarrhythmic | Ventricular tachycardia/fibrillation |
| Amitriptyline | Tricyclic Anti-depressant | Depression |
| Amlodipine | Calcium Channel Blocker | Hypertension |
| Amodiaquine | Anti-malaria | Malaria |
| Anakinra | IL-1R antagonist | RA, NOMID, CAPS |
| Anagrelide | PDE3 inhibitor (Platelet-reducing agent) | Essential thrombocythemia |
| Aprepitant | Substance P/neurokinin 1 (NK1) receptor antagonist | Nausea, vomiting |
| Aprotinin | Bovine pancreatic trypsin inhibitor (BPTI) | Perioperative blood loss |
| Arginine | Essential amino acid | Nutraceutical |
| Aripiprazole | Atypical antipsychotic | Bipolar disorder, major depressive disorder, autistic disorder |
| Artesunate | Antiprotozoal, antimalarian | Malaria |
| Atenolol | Competitive beta(1)-selective adrenergic antagonist | Hypertension, angina pectoris |
| Atorvastatin | Selective, competitive HMG-CoA reductase inhibitor (anti-cholesterol) | Coronary heart disease, acute coronary syndrome |
| Atovaquone | Synthetic hydroxynaphthoquinone (antiprotozoal, antimalarian) | Pneumocystis carinii pneumonia, toxoplasmosis |
| Atrial Natriuretic Peptide | Polypeptide vasodilator | Heart failure |
| Aspirin | NSAID | Pain, swelling, reduces the risk of a blood clot, prevent further heart attacks or strokes |
| Auranofin | Gold salt (anti-rheumatic) | RA |
| Azithromycin | Macrolide antibiotic, semi-synthetic | Bacterial infection, CAP, PID |
| Bazedoxifene | Selective estrogen receptor modulator | Osteoporosis |
| Bedaquiline | ATP-synthase inhibitor (antibiotic, diaryl-quinoline ) | Tuberculosis |
| Bemiparin | LMWH (anti-coagulant) | Venous thromboembolism, myocardial infarction |
| Benserazide | Peripherally-acting aromatic L-amino acid decarboxylase (AADC) or DOPA decarboxylase inhibitor | Parkinson's Disease |
| Bepridil | Calcium Channel Blocker | Hypertension and chronic stable angina |
| Bezafibrate | Fibrate | Hyperlipidemia |
| Biperiden | Muscarinic antagonist | Parkinson's Disease |
| Bosentan | Endothelin receptor antagonists | PAH |
| Bromocriptine | Dopamine agonist | Parkinson's Disease, prevention of lactation |
| Cabergoline | Dopamine receptor agonist | Hyperprolactinemia |
| Caffeine | CNS Stimulant | Newborn apnea |
| Calcitriol | Vitamin D3 | Vitamin |
| Candesartan | Angiotensin Receptor Blocker | Hypertension |
| Captopril | Angiotensin-converting-enzyme inhibitor | Anti-hypertension |
| Carbimazole | Imidazole-derivative (antithyroid) | Hyperthyroidism |
| Carglumic acid | Metabolic alkalosis agent | Hyperammonaemia in N-acetylglutamate synthase deficiency |
| Carvedilol | Betablocker | Hypertension |
| Celecoxib | NSAID (COX-2) | OA, RA, JRA, AS, acute pain, primary dysmenorrhea |
| Cephalexin | Cephalosporin antibiotic | Bacterial infections |
| Cholecalciferol | Vitamin D3 | Vitamin |
| Chlorpromazine | Phenothiazine (typical antipsychotic) | Psychotic disorders, nausea and vomiting, anxiety, hiccups |
| Chloroquine | Antimalarial and amebicidal drug | Malaria, Extraintestinal Amebiasis |
| Cidofovir | Deoxycytidine monophosphate analog | CMV-retinitis in AIDS |
| Cilnidipine | Calcium Channel Blocker | Hypertension |
| Cimetidine | Histamine H2-receptor blocker | Duodenal/gastric ulcers, GERD, pathological hypersecretory conditions |
| Ciprofloxacin | Fluoroquinolone | Antibiotic |
| Citalopram | Selective Serotonin Reuptake Inhibitors | Depression |
| Clarithromycin | Macrolide antibiotic | Bacterial infections |
| Clofoctol | Other antibiotic | Bacterial infections |
| Clomifene | Synthetic ovulation stimulant | Ovulatory dysfunction |
| Clomipramine | Tricyclic anti-depressant | Obsessive Compulsive Disorder |
| Clotrimazole | Imidazole derivative | Fungal infections |
| Colchicine | Antimitotic alkaloid | Gout |
| Dalteparin | LMWH (anti-coagulant) | DVT (prophylaxis), unstable angina/non-Q-wave myocardial infarction |
| Danazol | Antigonadotropins and similar agents | Endometriosis, fibrocystic breast disease, hereditary angioedema |
| Dapsone | Antibiotic | Dermatitis herpetiformis, leprosy |
| Deferoxamine | Iron chelating agent | Acute iron intoxication, chronic iron overload |
| Desmopressin | Vasopressin analogue | Diabetes Insipidus, bedwetting, hemophilia A, von Willebrand's disease |
| Diclofenac | NSAID | OA, RA, AS |
| Diflunisal | NSAID | OA, RA, mild to moderate pain |
| Digitoxin | Cardiac glycoside | Congestive HF, atrial fibrillation, atrial flutter, PAT, cardiogenic shock |
| Digoxin | Cardiac glycoside | Heart failure, atrial fibrillation |
| Dimethyl Fumarate | NA | Psoriasis, Multiple Sclerosis |
| Dipyridamole | Platelet aggregation inhibitor | Thromboembolism Prophylaxis Post-Cardiac Valve Replacement |
| Disulfiram | Alcohol antagonist | Chronic alcoholism |
| Donepezil | Acetylcholinesterase inhibitor | Alzheimer's Disease |
| Doxazosin |  | Anti-hypertensive |
| Doxycycline |  | Antibiotic |
| Ebastine | Histamine H1-receptor blocker | Allergies |
| Efavirenz |  | Anti-retroviral |
| Eflornithine |  |  |
| Enalapril | Angiotensin-converting-enzyme inhibitor | Anti-hypertensive |
| Enoxaparin | LMWH | Anti-coagulant |
| Epalrestat | Aldose reductase inhibitor | Anti-diabetic |
| Esomeprazole | Proton Pump Inhibitor | Antacid |
| Ethacrynic acid | Loop diuretic | Diuretic |
| Etodolac | NSAID (COX-2) | NSAID |
| Famotidine | Histamine H2-receptor blocker | Antacid |
| Fasudil |  | Vasodilator |
| Felodipine | Calcium Channel Blocker | Anti-hypertensive |
| Fenofibrate | Fibrate | Anti-cholesterol |
| Fingolimod | Sphingosine-1-phosphate receptor modulator | Multiple Sclerosis |
| Fish oil (EPA/DHA) |  | Anti-cholesterol |
| Flubendazole |  | Anti-parasitic |
| Fluoxetine |  | Anti-depressant |
| Fluspirilene |  | Anti-psychotic |
| Fluvastatin |  | Anti-cholesterol |
| Fluvoxamine | Serotonin antagonist and reuptake inhibitors | Anti-depressant |
| Glipizide | Sulfonylureas | Anti-diabetic |
| Glutamine |  | Nutraceutical |
| Griseofulvin |  | Anti-fungal |
| Haloperidol |  | Dopamine antagonist |
| Hydralazine |  | Anti-hypertensive |
| Hydroxychloroquine |  |  |
| Hymecromone |  | Antispasmodic |
| Ibandronate |  | Bisphosphonate |
| Ibuprofen |  | Analgesic |
| Imipramine | Tricyclic Anti-depressant | Anti-depressant |
| Imiquimod |  |  |
| Indomethacin |  | NSAID |
| Irbesartan | Angiotensin Receptor Blocker | Anti-hypertensive |
| Itraconazole |  | Anti-fungal |
| Ivermectin |  | Anti-parasitic |
| Ketoconazole |  | Anti-fungal |
| Ketorolac |  | NSAID |
| Lanreotide | Somatostatin analogue |  |
| Lansoprazole | Proton Pump Inhibitor | Antacid |
| Leflunomide |  | Arthritis |
| Levetiracetam |  | Anti-epileptic |
| Levofloxacin | Fluoroquinolone | Antibiotic |
| Licofelone |  | Osteoarthritis |
| Lithium |  | Bipolar disorders |
| Lidocaine |  | Anesthetic |
| Loperamide |  | Anti-diarrhea |
| Loratadine | Histamine H1-receptor blocker | Allergies |
| Losartan | Angiotensin Receptor Blocker | Anti-hypertensive |
| Lovastatin | Statin | Anti-cholesterol |
| Loxoprofen | NSAID | Anti-inflammatory |
| Macitentan | Endothelin receptor antagonists | Pulmonary arterial hypertension |
| Manidipine | Calcium Channel Blocker | Anti-hypertensive |
| Maraviroc | CCR5 receptor antagonist | Anti-retroviral |
| Mebendazole |  | Anti-parasitic |
| Meclofenamate |  |  |
| Megestrol acetate |  | Hormone |
| Mefloquine |  | Anti-malarial |
| Melatonin |  | Anti-insomnia |
| Meloxicam | NSAID | Anti-inflammatory |
| Memantine | NMDA receptor antagonist | Alzheimer's Disease |
| Mepacrine (Quinacrine) |  | Anti-parasitic |
| Metformin |  | Anti-diabetic |
| Methimazole |  |  |
| Methazolamide |  | Antiglaucoma, diuretic |
| Methylnaltrexone | μ-opioid antagonist | Opioid-induced constipation |
| Metoclopramide |  | Anti-emesis |
| Mifepristone |  | Abortifacient |
| Minocycline |  | Antibiotic |
| Mirtazapine |  |  |
| Montelukast |  |  |
| Mycophenolate |  | Immunosuppressant |
| Nadroparin |  |  |
| Naproxen |  | NSAID |
| Naltrexone |  | Opioid receptor antagonist |
| Nelfinavir |  | Anti-retroviral |
| Niclosamide |  | Anti-parasitic |
| Nifedipine | Calcium Channel Blocker | Anti-hypertensive |
| Nifurtimox |  |  |
| Nimodipine | Calcium Channel Blocker | Anti-hypertensive |
| Nisodipine | Calcium Channel Blocker | Anti-hypertensive |
| Nitazoxanide |  | Anti-protozoal |
| Nitisinone |  |  |
| Nitroxoline |  | Antibiotic |
| Nitroglycerine |  | Nitro-vasodilator |
| Norethandrolone | Androgen | Aplastic anemia |
| Noscapine |  | Anti-tussive |
| Octreotide |  |  |
| Olanzapine | Atypical antipsychotic | Anti-psychotic |
| Olsalazine |  | Anti-inflammatory |
| Omeprazole | Proton Pump Inhibitor | Antacid |
| Orlistat | Lipase Inhibitor | Obesity |
| Ormeloxifene | Selective estrogen receptor modulator | Contraceptive |
| Oseltamivir | Neuraminidase inhibitor | Anti-viral |
| Ouabain |  | Anti-aryhtmic |
| Oxcarbazepine | Voltage-gated sodium channel blocker | Epilepsy |
| Pantoprazole | Proton Pump Inhibitor | Antacid |
| Penfluridol | Diphenylbutylpiperidine | Anti-psychotic |
| Pentamidine |  | Anti-parasitic |
| Pentoxifylline | Xanthine derivative | Peripheral artery disease |
| Perphenazine | Phenothiazine | Anti-psychotic |
| Phenylbutyrate |  | Urea cycle disorders |
| Phentolamine | Alfa-adrenergic antagonist | Vasodilator |
| Phenytoin |  | Anti-epileptic |
| Pirfenidone |  | Anti-fibrotic |
| Pimozide | Diphenylbutylpiperidine | Anti-psychotic |
| Pioglitazone |  | Anti-diabetic |
| Plerixafor |  | Autologous HSCT |
| Pravastatin | Statin | Anti-cholesterol |
| Prazosin |  | Anti-hypertensive |
| Pregabalin | Gabapentinoid | Anti-convulsant |
| Promethazine | Phenothiazine | Anti-psychotic |
| Propranolol | Betablocker | Anti-hypertensive |
| Pyrimethamine | Dihydrofolate reductase inhibitor | Anti-parasitic |
| Pyrvinium pamoate |  | Anti-parasitic |
| Quetiapine | Atypical antipsychotic | Anti-psychotic |
| Rabeprazole | Proton Pump Inhibitor | Antacid |
| Ranolazine | Voltage-gated sodium channel blocker | Anti-angina |
| Repaglinide |  | Anti-diabetic |
| Ribavirin | nucleoside inhibitor | Anti-viral |
| Rifabutin |  | Antibiotic |
| Riluzole |  | ALS |
| Risperidone | Atypical antipsychotic | Anti-psychotic |
| Ritonavir | Protease inhibitor | Anti-HIV |
| Roflumilast | PDE-4 inhibitor | COPD |
| Rosuvastatin | Statin | Anti-cholesterol |
| Sertraline |  | Anti-depressant |
| Sildenafil | PDE-5 inhibitor | Erectile dysfunction |
| Simvastatin | Statin | Anti-cholesterol |
| Sirolimus |  | Inhibit organ transplant rejection |
| Sodium Bicarbonate |  | Antacid |
| Sodium Oxybate |  | Narcolepsy |
| Spironolactone |  | Anti-hypertensive |
| Sulfasalazine |  | Anti-rheumatic |
| Sulindac |  | NSAID |
| Tadalafil | PDE-5 inhibitor | Erectile dysfunction |
| Terbinafine |  | Anti-fungal |
| Telmisartan | Angiotensin Receptor Blocker | Anti-hypertensive |
| Tetrathiomolybdate |  | Copper toxicosis |
| Thalidomide | Immunomodulatory imide drug | Leprosy, multiple myeloma |
| Thiabendazole |  | Anti-parasitic |
| Thioridazine | Penothiazine | Anti-psychotic |
| Ticagrelor |  | Anti-platelet |
| Ticlopidine |  | Anti-platelet |
| Tigecycline | Glycylcycline | Antibiotic |
| Timolol | Betablocker | Anti-hypertensive |
| Tinzaparin | LMWH | Anti-coagulant |
| Tolfenamic acid | Fenamate NSAIDs | NSAID |
| Topiramate |  | Epilepsy |
| Tranexamic acid |  | Blood loss |
| Trazodone | Serotonin antagonist and reuptake inhibitors | Anti-depressant |
| Triamterene | Potassium-sparing diuretic | Diuretic |
| Trifluoperazine | Phenothiazine | Anti-psychotic |
| Ulinastatin | Trypsin inhibitor | Severe sepsis & pancreatitis |
| Urokinase |  |  |
| Valproic acid |  | Anti-convulsant |
| Valsartan | Angiotensin Receptor Blocker | Anti-hypertensive |
| Verapamil |  | Anti-hypertensive |
| Warfarin | Anti-Vitamin-K | Anti-coagulant |
| Zoledronate |  | Bisphosphonate |

**Table 2.** Some features of the 235 drugs listed.

|  | N | % |
| --- | --- | --- |
| **Human data** (*at least 1 case report, 1 obs. study or 1 clinical trial*) | **133** | **57%** |
| **At least 1 clinical trial** | **124** | **53%** |
| **Drug Off-Patent** | **176** | **75%** |

Several drugs have shown a survival benefit in randomized trials such as cimetidine (colorectal cancer) ^13^, progesterone (breast cancer) ^14^ or itraconazole (lung cancer)^15^. Of note, several other drugs induced responses in rare tumours, like clarithromycin ^16^, timolol ^17,18^ or propranolol ^19–22^.

## Discussion

We have found that the number of off-patent repurposing opportunities is large and increasing.

Until now, practice-changing examples have been limited despite the evidence. Joint non-commercial clinical development (academics, governments, charities) may bring new therapeutic options to patients at low cost, especially in indications for which the industry has no incentive to invest in. This may relieve healthcare systems currently under high financial stress.

A change in market authorisation regulation will be required to avoid off-label use. The Anticancer Fund is actively pursuing this innovative strategy, as exemplified by the granting of an EMA orphan designation for propranolol in angiosarcoma.

